# Two-photon excitation two-dimensional fluorescence spectroscopy (2PE-2DFS) of the fluorescent nucleobase 6-MI

**DOI:** 10.1101/2024.03.13.584863

**Authors:** Claire S. Albrecht, Lawrence F. Scatena, Peter H. von Hippel, Andrew H. Marcus

## Abstract

Base stacking is fundamentally important to the stability of double-stranded DNA. However, few experiments can directly probe the local conformations and conformational fluctuations of the DNA bases. Here we report a new spectroscopic approach to study the local conformations of DNA bases using the UV-absorbing fluorescent guanine analogue, 6-methyl isoxanthopterin (6-MI), which can be used as a site-specific probe to label DNA. In these experiments, we apply a two-photon excitation (2PE) approach to two-dimensional fluorescence spectroscopy (2DFS), which is a fluorescence-detected nonlinear Fourier transform spectroscopy. In 2DFS, a repeating sequence of four collinear laser pulses (with center wavelength ∼ 675 nm and relative phases swept at radio frequencies) is used to excite the lowest energy electronic-vibrational (vibronic) transitions of 6-MI (with center wavelength ∼ 340 nm). The ensuing low flux fluorescence is phase-synchronously detected at the level of individual photons and as a function of inter-pulse delay. The 2PE transition pathways that give rise to electronically excited state populations include optical coherences between electronic ground and excited states and non-resonant (one-photon-excited) virtual states. Our results indicate that 2PE-2DFS experiments can provide information about the electronic-vibrational spectrum of the 6-MI monomer, in addition to the conformation-dependent exciton coupling between adjacent 6-MI monomers within a (6-MI)_2_ dimer. In principle, this approach can be used to determine the local base-stacking conformations of (6-MI)_2_ dimer-substituted DNA constructs.

## 1. INTRODUCTION

The primary function of DNA is to encode (and protect) the genetic information as sequences of nucleobases, which may be ‘read’ and ‘manipulated’ by regulatory proteins that carry out the processes of genome maintenance and expression. DNA exists in its most stable form as the iconic Watson-Crick (WC) double-helix in which flanking bases are fully stacked and the sugar-phosphate backbones adopt a right-handed twist [1]. Nevertheless, under ‘physiological’ conditions the WC structure is only marginally stable. On local length scales the DNA duplex may undergo thermally induced fluctuations (termed DNA ‘breathing’) to populate local conformations in which flanking nucleobases are ‘twisted’ relative to their neighbors and thus somewhat more stacked or unstacked than in the canonical WC duplex. In such perturbed conformations, base-pairs are separated, and thus inter-base hydrogen bonds are broken, and/or local bases are fully unstacked and un-hydrogen-bonded and thus locally exposed to the external environment. Such unstable, non-canonical conformations may aid in the recognition, assembly and function of protein-DNA complexes involved in the core biological processes of DNA replication, transcription and repair [2].

Recent spectroscopic studies of DNA breathing in our laboratory have focused on (iCy3)_2_ dimer backbone-labeled single-stranded (ss) – double-stranded (ds) DNA fork constructs [3-7]. These experiments utilize the exciton-coupling between closely spaced monomers of the visible cyanine dye Cy3, which are attached ‘internally’ (referred to as ‘iCy3’) within the DNA sugar-phosphate backbones [8, 9]. When two iCy3 probes are positioned directly across the WC bridge on opposite complementary strands, they can form an exciton-coupled dimer probe, (iCy3)_2_, whose spectroscopic properties are sensitive to the local conformations of the DNA bases and sugar-phosphate backbones immediately adjacent to the probe. Linear and nonlinear optical spectroscopic measurements of (iCy3)_2_ dimer-labeled DNA constructs, in combination with theoretical analyses, have provided key insights about the mean local conformations and conformational disorder of these systems [4-7]. A recent development is the application of polarization-sweep single-molecule fluorescence (PS-SMF) spectroscopy to these systems, which provides information about the Boltzmann-weighted distributions of conformational macrostates and the microsecond dynamics of state-to-state interconversion [3].

Although the above studies on (iCy3)_2_ dimer-labeled DNA constructs can be used to infer information about the local conformations of DNA bases and sugar-phosphate backbones, they do not provide direct information about local base stacking. An alternative approach is to use fluorescent base analogues, which can be site-specifically substituted for native bases within a DNA construct. 6-methyl isoxanthoptherin (6-MI) is a fluorescent analogue of guanine that can form complementary hydrogen bonds with cytosine [10, 11]. However, unlike guanine, 6-MI absorbs at lower energy than any of the native DNA bases (at ∼340 nm) and emits fluorescence (at ∼430 nm) with relatively high quantum yield (∼0.7) [12]. In principle, spectroscopic studies of fluorescent mononucleotide and dinucleotide analogue-substituted DNA constructs can provide both direct information about the local conformations of the 6-MI dimer probes themselves, and also about the conformations of DNA bases immediately adjacent to the base analogue probe(s) [13, 14].

Two-dimensional fluorescence spectroscopy (2DFS) is a fluorescence-detected Fourier transform (FT) optical method that is well suited to measure the electronic-vibrational structure of fluorescent molecular chromophores, such as 6-MI [5, 7, 15, 16]. Like transmission-based multi-dimensional FT optical spectroscopies in the visible and IR regimes [17-19], 2DFS can reveal information about the optical transition pathways that are operative within a multi-level quantum molecular system. By virtue of its ability to sensitively monitor fluorescence against a dark background, 2DFS can detect relatively weak excited state population signals. Recent advances permit such phase-sensitive fluorescence-detected FT experiments to be carried out under low light signal conditions using single photon counting techniques [20].

In this work, we introduce a new experimental approach to perform 2DFS, which is based on two-photon excitation (2PE) of the UV-absorbing fluorescent base analogue 6-MI. Like the prior 2DFS experiments carried out on (iCy3)_2_ dimer-labeled DNA constructs, similar experiments carried out on 6-MI substituted DNA constructs hold promise to provide direct information about local base conformation and conformational distributions. In prior work by Widom *et al*., 2DFS was used to measure the exciton coupling of 2-aminopurine (2-AP) dinucleotide in solution – 2-AP is a fluorescent analogue of adenine [21]. Unfortunately, such experiments on UV-absorbing chromophores are challenging due to the technical difficulties of maintaining well-behaved broadband laser pulses in the UV spectral regime and detecting weak fluorescent signals above the background of scattered excitation light. In the following, we circumvent these difficulties by using a broadband pulsed laser source in the visible regime, with wavelength centered at ∼675 nm, to carry out 2PE-2DFS experiments.

Two-photon excitation (2PE) occurs when the energy of the source laser is one half of that of the lowest energy transition between ground state, |*g*⟩, and excited (final) state, |*f*⟩. While one-photon excitation (1PE) of the |*g*⟩ *⟶* |*f*⟩ transition is not dipole-allowed, 2PE can be achieved through a perturbative sequence of density-matrix elements that include coherences between |*g*⟩, intermediate ‘virtual’ states, |*e*⟩ and |*e*′⟩, and final states |*f*⟩ and |*f*′⟩ [22]. Here, the virtual states are assumed to have energies equal to one half of those of the excited states. Although the optical transitions involving virtual states do not absorb energy from the field, and thus do not lead to fluorescence, the virtual states do participate in coherences, e.g., |*g*⟩⟨*e*|, |*e*⟩⟨*f*| or |*e*⟩⟨*e*′|. The 2PE phenomenon was first proposed by Maria Goeppert Mayer in 1931 and demonstrated by Kaiser and Garrett in 1961 [23]. Because 2PE permits access to energy regimes previously difficult to study by 1PE, it has been useful to study the absorbance of fluorescent DNA base analogues [24].

Here we demonstrate 2PE-2DFS experiments performed on 6-MI monomer and dimer in buffered salt solution containing 100 mM NaCl, 6 mM MgCl_2_, and 10 mM TRIS at pH 7. These experiments employ ∼30 fs pulses with center wavelength ∼675 nm to excite electronic-vibrational transitions of the 6-MI chromophore, which has its peak absorbance at ∼340 nm. Our study reveals the presence of electronic-vibrational (vibronically) coupled transitions in 6-MI, which are otherwise difficult to observe in UV-visible linear absorbance spectra. These experiments serve as a proof-of-concept for future studies of the local conformations and conformation disorder of nucleic acid bases in DNA when subject to varying environmental conditions. The relatively high signal count rates we observe in our current experiments (∼100 kHz for ∼4 μM 6-MI in buffer solution) suggest that a variation of this approach can be extended to carry out measurements at the single-molecule level. Such experiments could, in principle, be applied to the direct study of the distributions and kinetics of base stacking conformational fluctuations, much like our group has done using the (iCy3)_2_ dimer-labeled DNA backbone probes [3].

## 2. EXPERIMENTAL METHODS

The 2PE-2DFS method employs similar principles to those of 1PE-2DFS [5, 15, 16, 21], in which the sample is illuminated by a repeating sequence of four collinear ultrafast laser pulses (see Fig. 1*A*). In our 2PE-2DFS experiments the pulse repetition rate is 250 kHz, the center wavelength is ∼675 nm, and the pulse full-width half-max (FWHM) bandwidth is ∼30 nm. Here the photon energy has been set to approximately one-half of the lowest energy transition of the 6-MI molecule (with peak absorption wavelength ∼340 nm). The pulses are characterized by their center times, *t*_*i*_, phases, *φ*_*i*_, inter-pulse delays, *t*_*ij*_ = *t*_*i*_ − *t*_*j*_, and relative phases, *φ*_*ij*_ [*i, j* ∈ {1,2,3,4}]. For a given measurement, the time delays *t*_21_, *t*_32_ and *t*_43_ are held fixed while the relative phases are swept continuously at tens-of-kilohertz frequencies.

**Figure 1.**
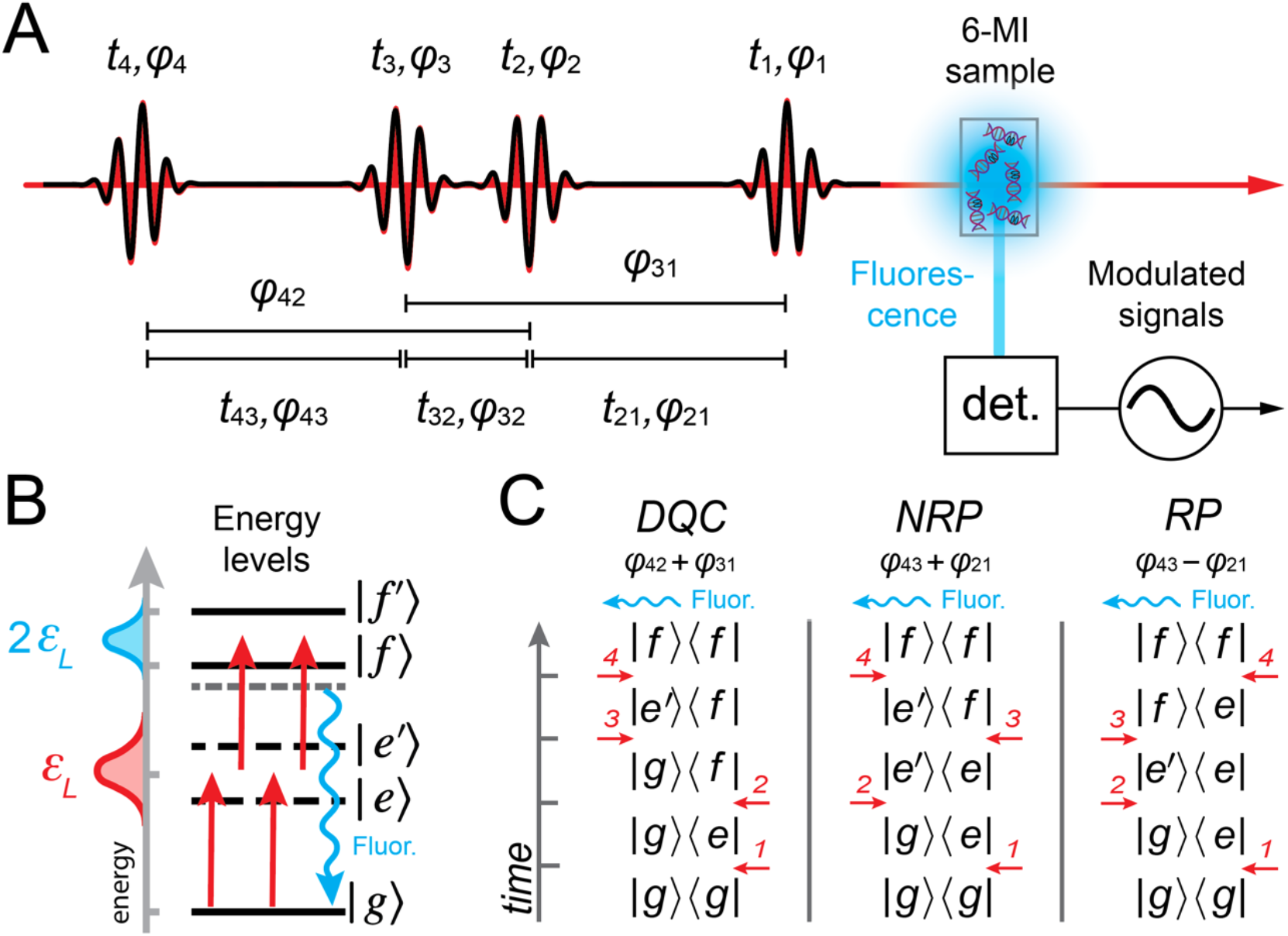
(***A***) The 2PE-2DFS method employs a repeating sequence of four collinear ultrafast laser pulses, which excites the 6-MI sample. Fluorescence from the sample is detected using phase sensitive methods. The relevant times and phases are indicated. (***B***) Energy level diagram indicating the laser pulse energy and molecular state energies. Fluorescence is spectrally separated from scattered laser excitation light. (***C***) Double-sided Feynman diagrams for 2PE-2DFS: DQC (double-quantum coherence, *φ*_42_ + *φ*_31_), NRP (non-rephasing, *φ*_43_ + *φ*_21_), and RP (rephasing, *φ*_43_ − *φ*_21_).

The four-pulse sequence induces one-photon transitions between ground state |*g*⟩, intermediate (‘virtual’) states, |*e*⟩ and |*e*′⟩, and final excited states |*f*⟩ and |*f*′⟩ (see Fig. 1*B*) [22]. The ensuing 2PE fluorescence signals are proportional to the populations of the excited states that vary at distinct radio frequencies, which can be detected separately using phase sensitive methods [20]. In Fig. 1*C* double-sided Feynman diagrams are shown, which indicate the three possible pathways of perturbative fourth-order density matrix elements that generate excited state populations. Each of the three Feynman pathways depends on a specific sequence of field-matter interactions that give rise to a unique phase-modulation signature. The ‘double-quantum coherence (DQC)’ pathway includes the intermediate coherence term |*g*⟩⟨*f*| between ground and excited states during the period *t*_32_, and has phase signature *φ*_42_ + *φ*_31_. In contrast, the ‘non-rephasing’ (NRP) and ‘rephasing’ (RP) pathways, which have phase signatures *φ*_43_ + *φ*_21_ and *φ*_43_ − *φ*_21_, respectively, include the intermediate coherence term |*e*′⟩⟨*e*| between distinct virtual states during the period *t*_32_. The recorded signals are complex-valued response functions of the inter-pulse delays: *S*_*DQC*_(*t*_21_, *t*_32_, *t*_43_), *S*_*NRP*_(*t*_21_, *t*_32_, *t*_43_) and *S*_*RP*_(*t*_21_, *t*_32_, *t*_43_). Fourier transformation of the response functions with respect to the delay variables yields the frequency- and phase-dependent 2D spectra: *Ŝ*_*DQC*_(*ω*_21_, *ω*_32_, *ω*_43_), *Ŝ*_*NRP*_(*ω*_21_, *ω*_32_, *ω*_43_) and *Ŝ*_*RP*_(*ω*_21_, *ω*_32_, *ω*_43_) [15, 16, 22].

In Fig. 2 is shown a schematic of the 2PE-2DFS instrument used in our experiments. The source laser is a custom-built, noncollinear optical parametric amplifier (NOPA), which is pumped by the pulsed output (250 kHz, 1039 nm, 6W) of a commercial ultrafast laser system (Pharos, Altos Photonics, Inc.). The NOPA produces tunable ultrafast pulses with power ∼300 mW, center wavelength ∼675 nm, pulse FWHM bandwidth ∼30 nm and compressed temporal FWHM duration of ∼33 fs (giving a time bandwidth product, ~ Δτ_*L*_(Δλ_*L*_*c*/λ^2^) ≈ 0.65, which is about 48% greater than the ideal value 0.44). The four-pulse sequence is generated using an arrangement of nested Mach-Zehnder interferometers (MZIs, see Fig. 2*A*). The NOPA output is initially passed through a single-prism pulse compressor for pre-dispersion compensation before it is split into two paths, each containing an equivalent MZI (labeled *A* and *B*). Detail of MZI *A* is shown in Fig. 2*B*. The MZI splits the pulse into two paths, which contain acousto-optic modulators (AOMs, Quanta-Tech, Inc.) that are swept continuously at MHz frequencies. The driving frequencies of the AOMs are detuned so that a time-dependent relative phase shift is imparted to successive pulse-pairs traveling through the MZI: *φ*_*ij*_ = *φ*_*j*_ − *φ*_*i*_ = 2π*v*_*ij*_*t*. For NRP and RP measurements, MZI *A* has *v*_21_ = 5 kHz and MZI *B* has *v*_43_ = 8 kHz. When measuring the DQC signal the modulation frequencies are set such that *v*_31_ = 5 kHz and *v*_42_ = 8 kHz. The inter-pulse delays, *t*_*ij*_, are controlled using nano-positioning translation stages (Aerotech, Inc.). In a typical data run, the stages were stepped in 4 fs increments over a full range of ∼150 fs.

**Figure 2.**
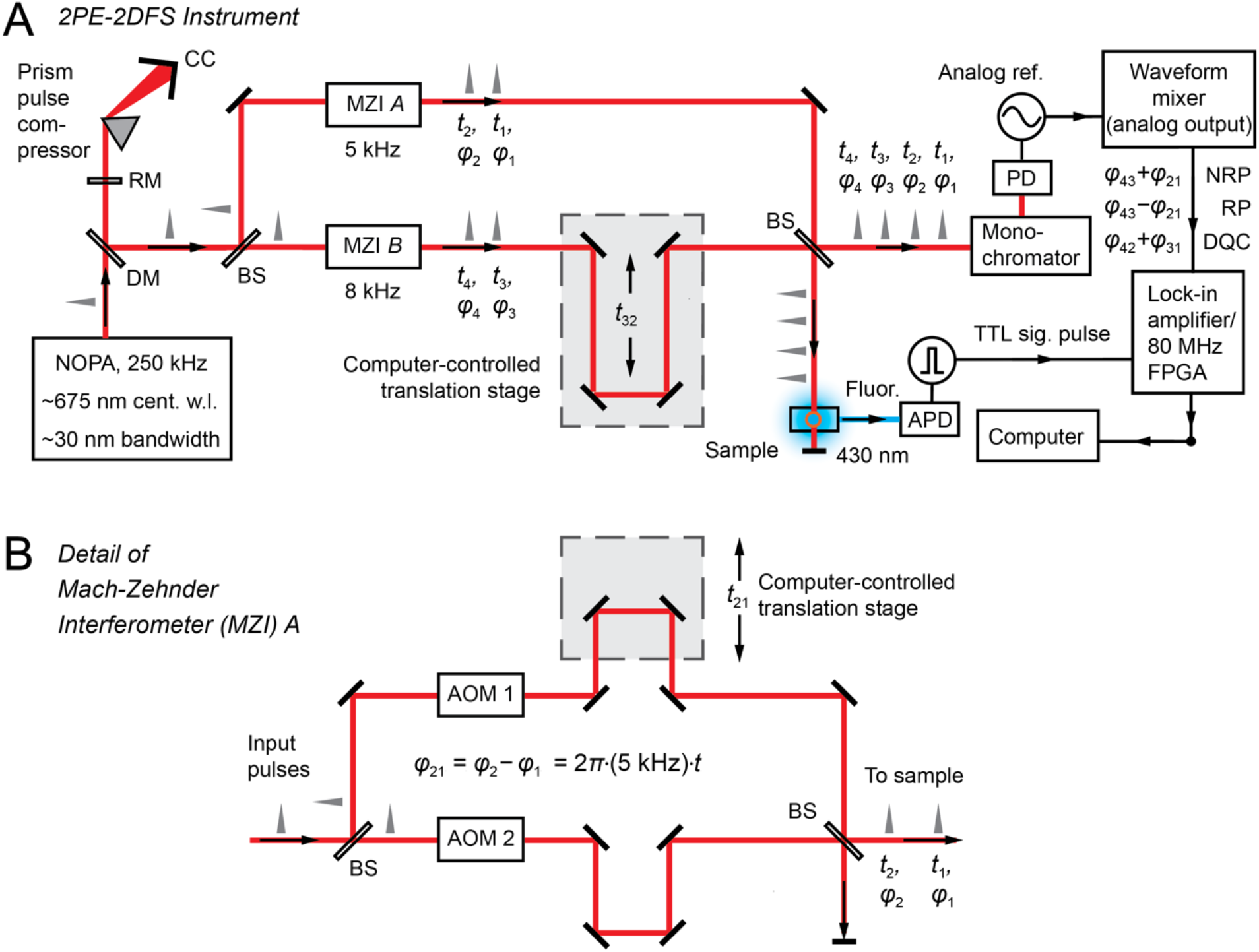
(***A***) Schematic of the 2PE-2DFS experimental setup. (***B***) Detail of the Mach-Zehnder interferometer (MZI) *A*. See text for discussion. NOPA: non-collinear optical parametric amplifier; CC: corner-cube; DM: d-shaped mirror; BS: beam-splitter; APD: avalanche photodiode; PD: photodiode; TTL: transistor-transistor logic.

The pulse sequence that exits the final recombining beam splitter is incident at the 6-MI sample with ∼10 mW of power. Individual fluorescent photons are detected at 90° using a 5 cm collection lens and a fiber-coupled avalanche photodiode (APD, Laser Components, COUNT-10B-FC, dark count ∼10 Hz). This detection geometry was chosen to minimize the amount of scattered excitation light that entered the detection channel. A replica copy of the four-pulse sequence is used to construct reference waveforms for the phase-sensitive detection electronics. The reference beam is directed through a monochromator and detected using a photodiode. The fundamental phase shifts (*φ*_43_, *φ*_21_, *φ*_42_, *φ*_31_) are converted into digital waveforms and recorded by a field programmable gate array (FPGA) for low-signal-flux measurements (see below). Alternatively, these waveforms are mixed to generate the analog sum frequency waveforms, *φ*_43_ + *φ*_21_ and *φ*_42_ + *φ*_31_ (for detecting NRP and DQC signals, respectively) and the difference frequency waveform, *φ*_43_ − *φ*_21_ (for detecting the RP signal).

Depending on the (sample-specific) fluorescence intensity, we performed our experiments in either the ‘high-flux’ (≳ 100,000 cps) or ‘low-flux’ (≲ 100,000 cps) signal regimes. In the high-flux regime, the analog reference waveforms were used to trigger a lock-in amplifier, which integrates the fluorescence over many phase-modulation cycle periods (lock-in time constant ∼2 ms) and isolates the real and imaginary parts of complex-valued response functions, *S*_*DQC*_, *S*_*NRP*_ and *S*_*RP*_ [15, 16]. In the low-flux signal regime, the fundamental reference waveforms are sent to an FPGA, which records and stores the values of the phase shifts for each single-photon detection event (clock speed 80 MHz) using the method of phase-tagged photon counting (PTPC) [20]. In post-data-acquisition, the discrete PTPC signals are integrated over a fixed bin period (∼2 ms) to construct the complex-valued response functions following a procedure outlined in prior work [3, 20].

## 3. RESULTS AND DISCUSSION

We performed our 2PE-2DFS measurements on solutions containing 6-MI mononucleoside (MNS) and, in separate experiments, (6-MI)_2_ dinucleotide di-phosphate (DNTDP). We prepared these samples in a 5 mm quartz cuvette at 4 *μ*M concentration in standard aqueous buffer salt solution containing 10 mM TRIS (pH 7.0), 100 mM NaCl and 6 mM MgCl_2_. As mentioned previously, we tuned the excitation source (675 nm) to approximately one half the energy of the lowest energy transition (∼ 340 nm) in the 6-MI molecule. In Figs. 3*A* and 3*B* we show the molecular structures of the 6-MI MNS and the (6-MI)_2_ DNTDP, respectively. The electric transition dipole moment (EDTM) of the lowest energy transition (*S*_1_) is indicated as a double-headed blue arrow, which lies within the plane of the 6-MI nucleobase [25]. The *S*_1_transition is coupled to at least one effective vibrational mode with energy, *ħω*_*vib*_ ∼400 cm^-1^. If the two bases of the (6-MI)_2_DNTDP are closely spaced (e.g., if they are ‘stacked’), the component EDTMs can couple electrostatically (with Davydov splitting 2*J*) to produce coherent symmetric (+) and anti-symmetric (–) superpositions of electronic-vibrational (vibronic) products (i.e., molecular excitons, see Fig. 3*C*). The precise geometric arrangement of the coupled EDTMs will determine the dipole strengths and energies of the (6-MI)_2_ DNTDP spectral features.

**Figure 3.**
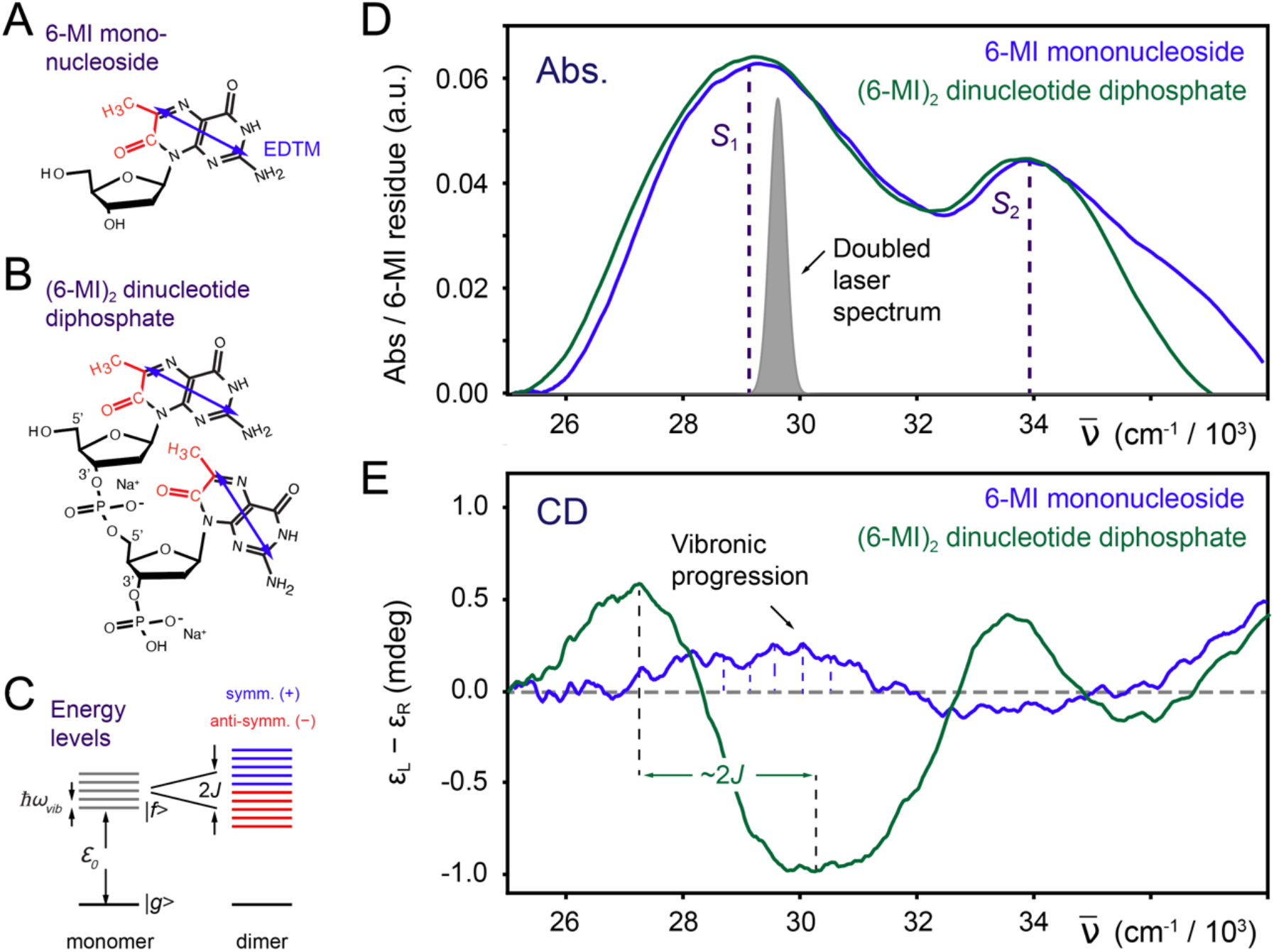
(***A***) Molecular structure of the 6-MI mononucleoside (MNS). Atoms that are additional to those of the natural base guanine are indicated in red. The blue double-headed arrow indicates the direction of the lowest energy electric dipole transition moment (EDTM). (***B***) Structure of the (6-MI)_2_ dinucleotide diphosphate (DNTDP). (***C***) Energy level diagram of the 6-MI MNS (monomer) and the (6-MI)_2_ DNTDP (dimer) in the presence of electrostatic coupling, *J*, between EDTMs. The monomer electronic transition with energy, *ε*_0_, is coupled to a vibrational mode with energy spacing, *ħω*_*vib*_. The coupling between monomers introduces symmetric (+, blue) and anti-symmetric (−, red) electronic-vibrational (vibronic) coherent states. (***D***) Absorbance spectra of the 6-MI MNS (blue) and the (6-MI)_2_ DNTDP (green) in aqueous buffer salt solution (10 mM TRIS, 100 mM NaCl, and 6 mM MgCl_2_). Vertical dashed lines indicate the lowest energy electronic transition (*S*_1_) at ∼29,300 cm^-1^ and the second lowest transition (*S*_2_) at 34,000 cm^-1^ of the 6-MI MNS. The doubled laser spectrum used in the 2PE-2DFS experiments is shown in gray. (***E***) Circular dichroism (CD) spectra of the 6-MI MNS (blue) and (6-MI)_2_ DNTDP (green).

In Figs. 3*D* and 3*E* we show the experimental absorbance and circular dichroism (CD) spectra, respectively, of the 6-MI MNS (blue) and the (6-MI)_2_ DNTDP (green). The peak absorbance of the 6-MI MNS *S*_1_ transition at ∼29,300 cm^-1^ (∼341 nm) is significantly lower than the peak absorbance of the second *S*_2_ transition at ∼34,000 cm^-1^ (∼294 nm). The *S*_1_ transition is relatively broad, and its underlying vibronic features are only barely perceptible. We note that Stanley and co-workers were able to resolve vibronic transitions of 6-MI using Stark spectroscopy [12]. The absorbance spectrum of the (6-MI)_2_ DNTDP exhibits additional broadening of the *S*_1_ line shape in comparison to that of the 6-MI MNS, which is expected due to the spectral shifts associated with the electrostatic coupling, *J*. Also shown in Fig. 3*D* is the frequency-doubled laser spectrum used in our 2PE-2DFS experiments. The vibronic fine structure can be more easily discerned in the CD spectra of the 6-MI MNS and the (6-MI)_2_ DNTDP, which are shown in Fig. 3*E*.

The CD spectrum is the difference in absorbance between left- and right-circularly polarized light and is sensitive to the chiral environment immediately surrounding the EDTM [26, 27]. If the direction of the EDTM depends on the nuclear coordinates of the 6-MI molecule (i.e., a violation of the Condon approximation and the presence of Hertzberg-Teller coupling), the vibronic transitions will give rise to optical activity apparent in the CD spectrum [28-30]. The CD of the 6-MI MNS exhibits a weakly positive band of vibronic features over the range ∼26,500 cm^-1^ to ∼32,000 cm^-1^, corresponding to the *S*_1_ transition, and a weakly negative band of vibronic features over the range ∼32,000 cm^-1^ to ∼33,000 cm^-1^, corresponding to the *S*_2_ transition. In contrast, the CD of the (6-MI)_2_ DNTDP exhibits a strongly positive band (peaked at ∼27,000 cm^-1^) and a strongly negative band (peaked at ∼30,000 cm^-1^), which symmetrically splits the *S*_1_ transition. Such behavior is characteristic of a right-handed Cotton effect that is due to presence of electrostatic coupling between the degenerate EDTMs of the component 6-MI monomers [26]. The peak-to-peak splitting is ∼2,800 cm^-1^, suggesting an electrostatic coupling *J* ≲ 1,400 cm^-1^. We note that a right-handed Cotton effect appears to be operative for the *S*_2_ transition, like the effect seen for the *S*_1_ transition.

In Fig. 4 we present the results of our 2PE-2DFS experiments carried out on the 6-MI MNS. In each panel the real part of the 2PE-2DFS spectrum is plotted for a specific phase condition (from left to right: DQC, NRP and RP) and pulse scanning condition, where one of the inter-pulse delays is set to zero (from top to bottom: *t*_32_ = 0, *t*_21_ = 0 and *t*_43_ = 0) while the remaining delay variables are scanned. The 2D spectra are constructed by taking the Fourier transforms of the measured response functions with respect to the scanned delay variables. The resulting 2D spectra are presented as contour diagrams as a function of the frequencies of the scanned delays. In the horizontal and vertical margins of the 2D spectra are shown the linear absorbance spectrum of 6-MI and the laser spectrum plotted at the two-photon energy.

**Figure 4.**
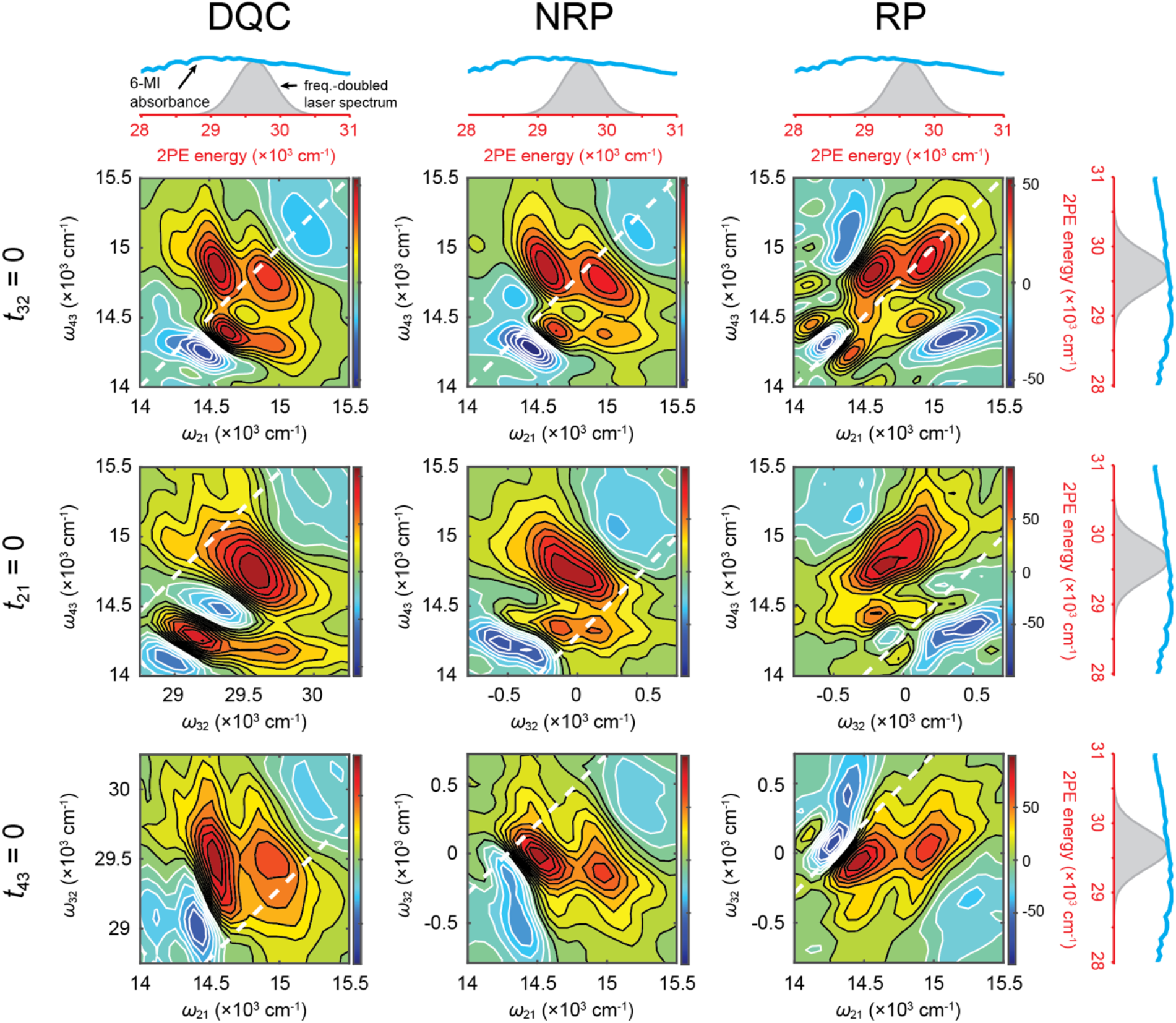
2PE-2DFS experimental results for the 6-MI MNS in aqueous buffer salt solution (10 mM TRIS, 100 mM NaCl, and 6 mM MgCl_2_). The real parts of the experimental spectra are plotted as 2D contour diagrams. Columns indicate a specific signal phase condition (from left to right: DQC, NRP and RP) and rows indicate one of the three inter-pulse delays set to zero (from top-to-bottom: *t*_*ji*_ = 0, *T*_21_ = 0 and *t*_43_ = 0). In the margins are shown the linear absorbance spectrum of the 6-MI MNS (blue) and the spectrum of the laser at twice its energy (gray).

We note that the 2D spectrum for each of the nine experimental conditions exhibits peaks and cross-peaks, which occur at energies consistent with the presence of vibronic transitions described by the double-sided Feynman diagrams shown in Fig. 1*C*. For all the 2PE-2DFS data, spectral features corresponding to vibronic fine structure can be clearly discerned. For example, all the 2PE-2DFS spectra for *t*_32_ = 0 (top row) exhibit four prominent features with spacing ∼400 cm^-1^, which are shifted vertically below the diagonal (along the *ω*_43_ axis) by ∼200 cm^-1^. These features occur at approximately one half the transition energies expected of vibronic features associated with the *S*_1_ transition, which can be seen in the CD spectrum shown in Fig. 3*E*. 2DFS experiments in which the intermediate states (|*e*⟩, |*e*′⟩) are one-photon resonant are often carried out with *t*_32_ = 0 and scanning the *t*_21_ and *t*_43_ delay variables. It is interesting to note that the general structures and symmetries of the 2PE-2DFS line shapes [*Ŝ*_*DQC*_(*ω*_21_, *t*_32_ = 0, *ω*_43_), *Ŝ*_*NRP*_(*ω*_21_, *t*_32_ = 0, *ω*_43_) and *Ŝ*_*RP*_(*ω*_21_, *t*_32_ = 0, *ω*_43_)] are essentially analogous to those of 1PE-2DFS, except that the spectral features occur at the one-photon transition energies at the frequencies *ω*_21_ and *ω*_43_ are associated with coherences between ground state and virtual states (e.g., |*g*⟩⟨*e*| and |*g*⟩⟨*e*′|) and between virtual states and final states (e.g., |*e*⟩⟨*f*′|, |*e*′⟩⟨*f*|). However, for the 2PE-2DFS spectra with *t*_21_ = *t*_43_ = 0 (second and third rows of Fig. 4, respectively), prominent spectral features for the DQC phase condition occur at two-photon transition energies at the frequency *ω*_32_ associated with coherences between ground state and final states (e.g., |*g*⟩⟨*f*′|, |*g*⟩⟨*f*|). This contrasts with the NRP and RP phase conditions in which spectral features occur at relatively low energies at the frequency *ω*_32_ associated with coherences between virtual vibronic states (e.g., |*e*⟩⟨*e*′|) with vibrational energy spacings. These results suggest that 2PE-2DFS experiments on the 6-MI MNS can be analyzed using a model Hamiltonian to extract quantitative information about the electronic-vibrational states of the chromophore in its monomeric form.

In Fig. 5 we present the results of our 2PE-2DFS experiments carried out on the (6-MI)_2_ DNTDP. These data show that the 2D spectral features of the (6-MI)_2_ DNTDP are generally narrower than those of the 6-MI MNS, and in some cases the spectral features appear to split into additional peaks. In particular, the 2D peak widths along the *ω*_32_ axes for *t*_21_ = *t*_43_ = 0 spectra appear to be most sensitive. It is possible that this apparent sensitivity is related to the absence of an obscurring population contribution to the 2PE-2DFS signal during the *t*_32_ interval (see Fig. 1*C*). The absence of population contributions to the NRP and RP signals in 2PE-2DFS is a unique situation in comparison to 1PE-2DFS, in which case the intermediate states are one-photon resonant and population contributions dominate. Our results for the (6-MI)_2_ DNTDP are consistent with the presence of conformation-dependent electrostatic coupling, *J*, between the 6-MI nucleobases within the (6-MI)_2_ DNTDP. These observations suggest that future studies that compare 2PE-2DFS data to theoretical models may be used to determine structural information about local base conformation and disorder in these and related systems.

**Figure 5.**
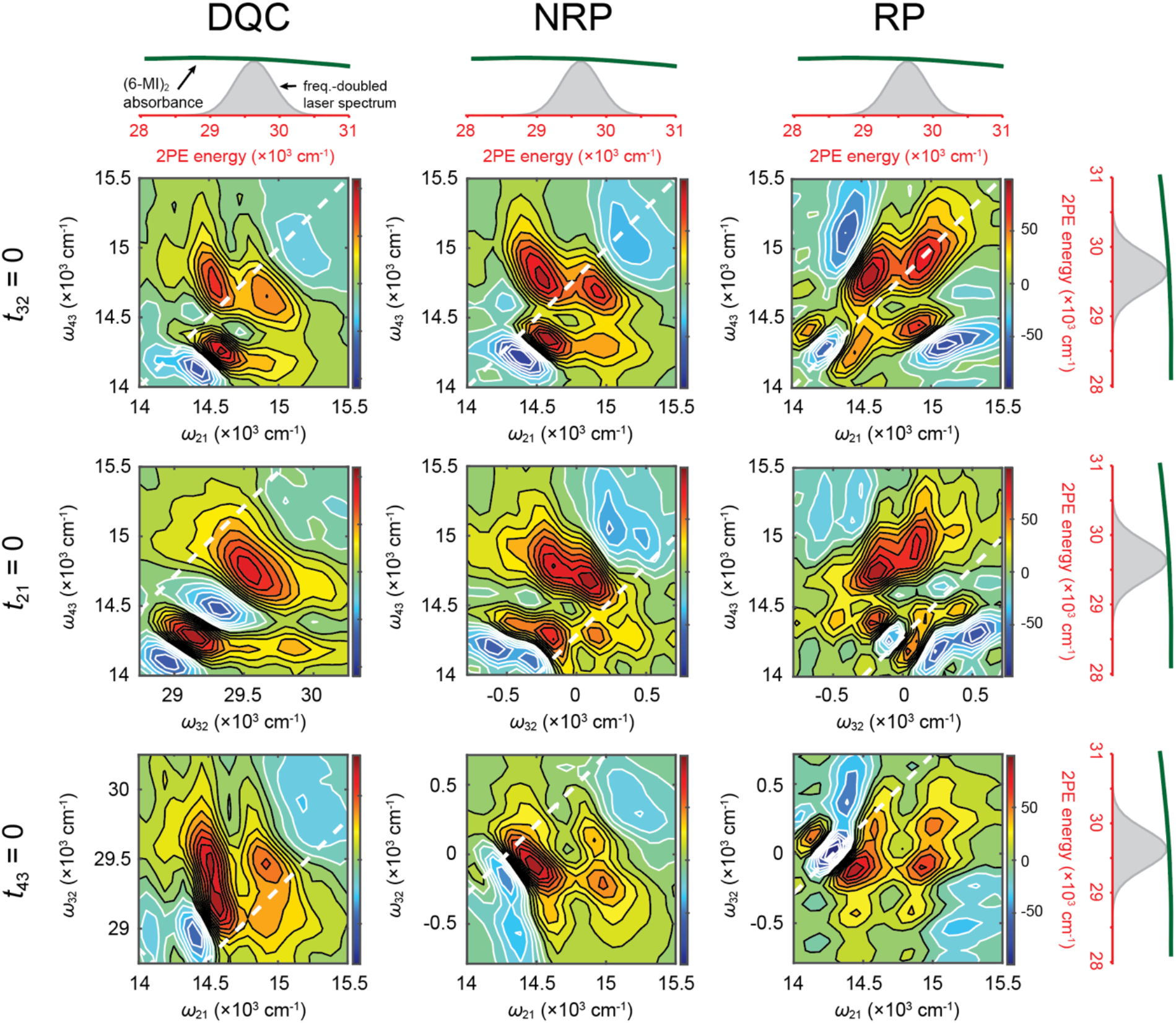
2PE-2DFS experimental results for the (6-MI)_2_ DNTDP in aqueous buffer salt solution (10 mM TRIS, 100 mM NaCl, and 6 mM MgCl_2_). See Fig. 4 caption for further explanation.

We emphasize that for both the 6-MI MNS and (6-MI)_2_ DNTDP, the 2PE-2DFS spectra exhibit detailed vibronic structure that is not well resolved in the UV-visible absorbance spectrum (Figs. 3*D*) and is only weakly resolved in the CD spectrum (Fig. 3*E*). These results suggest that 2PE-2DFS may be a generally useful approach to probe the vibronic structure of UV-absorbing molecules and molecular aggregates.

## 4. CONCLUSIONS

In this work, we have demonstrated a two-photon excitation approach to two-dimensional fluorescence spectroscopy (2PE-2DFS) to study the UV-absorbing fluorescent guanine analogue, 6-MI. The 2PE-2DFS spectra reveals electronic-vibrational (vibronic) transitions in 6-MI, which are otherwise difficult to resolve with standard spectroscopic methods such as UV-visible absorption and CD spectroscopy. The sensitivity of the vibronic features to local electrostatic coupling is evident from the differences between 2PE-2DFS spectra of 6-MI MNS versus (6-MI)_2_ DNTDP (compare Fig. 4 to Fig. 5). This sensitivity may be related to the lack of population contributions to the NRP and RP signals during the *t*_32_ interval (as illustrated in Fig. 1*C*), which is a unique advantage of the 2PE-2DFS approach.

In future studies, we will develop quantum mechanical models to simulate the absorbance, CD and 2PE-2DFS spectra of the 6-MI MNS to better understand the nature of the electronic-vibrational coupling. Extensions of these models to account for the electrostatic coupling between component nucleobases will permit us to simulate the 2PE-2DFS data that we have obtained for the (6-MI)_2_ DNTDP. While we have focused on the fluorescent base analogue 6-MI for its potential as a probe of local base stacking conformations in DNA, we note that the 2PE-2DFS method might be more generally applied to study electronic-vibrational structure of other UV-absorbing fluorescent chromophores.

The 2PE-2DFS experiments of the current work set the stage for future investigations of local base conformations of (6-MI)_2_ dinucleotide-substituted DNA constructs. Such experiments could reveal important information about local base conformations at specific positions in DNA, and its influence on protein-DNA interactions. It is interesting to note that the signal rates of our current ensemble 2PE measurements (∼100,000 cps) suggest the feasibility of performing similar measurements at the single-molecule level. Such an extensions of the 2PE-2DFS approach to the single-molecule level will, in principle, enable studies of the local fluctuations of DNA bases (i.e., DNA ‘breathing’) in duplex DNA constructs in real time.

## ACKNOWLEDGEMENTS

The authors thank their laboratory colleagues for engaging in many insightful discussions. This work was funded by the National Institute of Health General Medical Sciences (Grant No. GM-15792 to A.H.M. and P.v.H.). P.v.H. is an American Cancer Society Research Professor of Chemistry.

## REFERENCES

[1] R. R. Sinden, [DNA Structure and Function] Academic Press, San Diego (1994).

[2] P. H. von Hippel, N. P. Johnson, and A. H. Marcus, “50 years of DNA ‘breathing’: Reflections on old and new approaches,” Biopolymers, 99, 923–954 (2013).

[3] J. Maurer, C. S. Albrecht, P. Herbert, D. Heussman, A. Chang, P. H. von Hippel, and A. H. Marcus, “Studies of DNA ‘breathing’ by polarization-sweep single-molecule fluorescence microscopy of exciton-coupled (iCy3)2 dimer-labeled DNA fork constructs,” J. Phys. Chem. B, 127, 10730–10748 (2023).

[4] D. Heussman, L. Enkhbaatar, M. I. Sorour, K. A. Kistler, P. H. von Hippel, S. Matsika, and A. H. Marcus, “Using transition density models to interpret experimental optical spectra of exciton-coupled cyanine (iCy3)2 dimer probes of local DNA conformations at or near functional protein binding sites,” Nucl. Acids Res., (2023).

[5] D. Heussman, J. Kittell, P. H. von Hippel, and A. H. Marcus, “Temperature-dependent local conformations and conformational distributions of cyanine dimer labeled single-stranded–double-stranded DNA junctions by 2D fluorescence spectroscopy,” J. Chem. Phys., 156(4), (2022).

[6] D. Heussman, J. Kittell, L. Kringle, A. Tamimi, P. H. von Hippel, and A. H. Marcus, “Measuring local conformations and conformational disorder of (Cy3)2 dimers labeled DNA fork junctions using absorbance, circular dichroism and two-dimensional fluorescence spectroscopy,” Faraday Disc., 216, 211–235 (2019).

[7] L. Kringle, N. Sawaya, J. R. Widom, C. Adams, M. G. Raymer, A. Aspuru-Guzik, and A. H. Marcus, “Temperature-dependent conformations of exciton-coupled Cy3 dimers in double-stranded DNA,” J. Chem. Phys., 148, (2018).

[8] M. Levitus, and S. Ranjit, “Cyanine dyes in biophysical research: The photophysics of polymethine fluorescent dyes in biomolecular environments,” Quart. Revs. Biophys., 44(1), 123–151 (2011).

[9] W. Lee, P. H. von Hippel, and A. H. Marcus, “Internally labeled Cy3 / Cy5 DNA constructs show greatly enhanced photostability in single-molecule FRET experiments,” Nucl. Acids Res., 42, 5967–5977 (2014).

[10] M. E. Hawkins, “Fluorescent pteridine nucleoside analogs: a window on DNA interactions,” Cell biochemistry and biophysics, 34(2), 257–81 (2001).

[11] E. Seibert, A. S. Chin, W. Pfleiderer, M. E. Hawkins, W. Laws, R. Osman, and J. B. Ross, “pH-dependent spectroscopy and electronic structure of the guanine analogue 6,8-dimethylisoxanthopterin,” J. Phys. Chem. A, 107, 178–185 (2003).

[12] G. Kodali, M. Narayanan, and R. J. Stanley, “Excited-state electronic properties of 6-methylisoxanthopterin (6-MI): an experimental and theoretical study,” J. Phys. Chem. B, 116(9), 2981–2989 (2012).

[13] B. R. Camel, D. Jose, K. Meze, A. Dang, and P. H. von Hippel, “Mapping DNA conformations and interactions within the binding cleft of bacteriophage T4 single-stranded DNA binding protein (gp32) at single nucleotide resolution,” Nucl. Acids Res., 49, 916–927 (2021).

[14] D. Jose, S. E. Weitzel, W. A. Baase, and P. H. von Hippel, “Mapping the interactions of the single-stranded DNA binding protein of bacteriophage T4 (gp32) with DNA lattices at single nucleotide resolution: gp32 monomer binding,” Nucl. Acids Res., 43, 9276–9290 (2015).

[15] A. Perdomo, J. R. Widom, G. A. Lott, A. Aspuru-Guzik, and A. H. Marcus, “Conformation and electronic population transfer in membrane supported self-assembled porphyrin dimers by two-dimensional fluorescence spectroscopy,” J. Phys. Chem. B, 116, 10757–10770 (2012).

[16] P. F. Tekavec, G. A. Lott, and A. H. Marcus, “Fluorescence-detected two-dimensional electronic coherence spectroscopy by acousto-optic phase modulation,” J. Chem. Phys., 127, 214307 (2007).

[17] M. Reppert, and A. Tokmakoff, “Computational amide I 2D IR spectroscopy as a probe of protein structure and dynamics,” Ann. Rev. Phys. Chem., 67, 359–386 (2016).

[18] A. Ghosh, J. S. Ostrander, and M. T. Zanni, “Watching proteins wiggle: Mapping structures with two-dimensional infrared spectroscopy,” Chem. Revs., 117, 10726–10759 (2017).

[19] D. M. Jonas, “Optical analogs of 2D NMR,” Science, 300, 1515–1517 (2003).

[20] A. Tamimi, T. Landes, J. Lavoie, M. G. Raymer, and A. H. Marcus, “Fluorescence-detected Fourier transform electronic spectroscopy by phase-tagged photon counting,” Opt. Express, 28, 25194–25214 (2020).

[21] J. R. Widom, N. P. Johnson, P. H. von Hippel, and A. H. Marcus, “Solution conformation of 2-Aminopurine (2-AP) dinucleotide by ultraviolet 2D fluorescence spectroscopy (UV-2D FS),” New Journal of Physics, 15, 025028–43 (2013).

[22] M. G. Raymer, Landes, T., and A. H. Marcus, “Entangled two-photon absorption by atoms and molecules: A quantum optics tutorial,” J. Chem. Phys., 155, 081501-1-25 (2021).

[23] W. Kaiser, C. G. B. Garrett, “Two-photon excitation in CaF2:Eu2+,” Phys. Rev. Lett., 7, 229–231 (1961).

[24] A. Mikhaylov, S. de Reguarati, J. Pahapill, P. R. Callis, B. Kohler, and A. Rebane, “Two-photon absorption spectra of fluorescent isomorphic DNA base analogs,” Biomed. Optics Express, 9, 447–452 (2018).

[25] J. R. Widom, D. Rappoport, A. Perdomo-Ortiz, H. Thomsen, N. P. Johnson, P. H. von Hippel, A. Aspuru-Guzik, and A. H. Marcus, “Electronic transition moments of 6-methyl isoxanthoptherin (6-MI) - a fluorescent analog of guanine,” Nucl. Acids Res., 41, 995–1004 (2012).

[26] B. Nordén, A. Rodger, and T. Dafforn, [Linear Dicrhoism and Circular Dichroism: A Textbook on Polarized-Light Spectroscopy] RSC Publishing Cambridge, UK (2010).

[27] J. Michl, and E. W. Thulstrup, [Spectroscopy with polarized light: Solute alignment by photoselection, in liquid crystals, polymers and membranes] VCH Publishers, Inc., (1995).

[28] A. Baiardi, J. Bloino, and V. Barone, “General time dependent approach to vibronic spectroscopy including Franck-Condon, Hertzberg-Teller, and Duschinsky effects,” J. Chem. Theory Comput., 9, 4097–4115 (2013).

[29] S. Kundu, P. P. Roy, G. R. Fleming, and N. Makri, “Franck-Condon and Hertzberg-Teller signatures in molecular absorption and emission spectra,” J. Phys. Chem. B, 126, 2899–2911 (2022).

[30] M. Nooijen, “Investigation of Herzberg-Teller Franck-Condon approaches and classical simulations to include effects due to vibronic coupling in circular dichroism spectra: The case of dimethyloxirane continued,” Int. J. Quant. Chem., 106, 2489–2510 (2005).

